# Development of Shelf-Stable Reagents and Assay Kits for Bioluminescence Applications using the Capillary-Assisted Vitrification Platform Stabilization Technology

**DOI:** 10.64898/2026.07.11.737891

**Authors:** Shari M. Radford, Yolanda Peris-Taverner, Michael R. C. Dibble, Kevin C. Corn, Tian Zhu, Shannon E. Martello, McKenzie A. Mayeaux, Amanda Ladd, Sankar Renu, Tejasvi Chunduri, Aniket C. Jadhav, Melanie Dart, Marjan Rafat, Mary Shank-Retzlaff, Laura L. Bronsart

**Affiliations:** Ambient Biosciences, 1600 Huron Parkway, Bldg 520 Rm 2390, Ann Arbor, MI, USA; Promega Corp., 2800 Woods Hollow Rd. Madison, WI, USA; Department of Chemical and Biomolecular Engineering, Vanderbilt University, Nashville, TN, USA; Department of Biomedical Engineering, Vanderbilt University, Nashville, TN, USA; Department of Radiation Oncology, Vanderbilt University Medical Center, Nashville, TN, USA

**Author notes:** Corresponding Author: Mary Shank-Retzlaff.

**Keywords:** Bioluminescence, luciferase, reagent stability, luciferin

## Abstract

Luminescence is a powerful method for detecting trace analytes and monitoring biological processes. However, most bioluminescence reagents, including luciferase and its substrates, are sensitive to temperature, limiting their useable shelf lives, and resulting in inconsistent performance. Enhancing the stability of these reagents could improve data quality, simplify workflows, and address cold chain storage issues. In this study, we demonstrate the application of the platform stabilization technology, capillary-assisted vitrification (CAV), as a tool to stabilize different luciferases and their substrates, and the application of the stabilized reagents in both *in vitro* and *in vivo* bioluminescent assays. We demonstrate that CAV-stabilized reagents can be stored and shipped ambiently, maintain consistent performance over time, and are suitable for use in cell viability quantification, tumor monitoring, *in vivo* imaging, microbial detection, and immunoassays. Additionally, different reagents can be co-formulated to make ready-to-use assay kits that can also be shipped and stored ambiently. Our results demonstrate that CAV stabilization is a viable alternative to traditional storage methods, with broad potential to improve bioluminescence workflows.

## Introduction

Luminescence has emerged as a powerful tool for biological and biochemical research (1, 2, 3, 4, 5). Bioluminescence imaging is widely applied to identify and monitor tumor growth and to study viral infection in animal models (6, 7, 8). When combined with *in vivo* cell culture and reporter genes, luminescence can be used to assess both the efficacy and toxicity of drug candidates (9). Luminescence is also used for cell viability analysis and detection of microbial contamination in hospital and food production settings (10, 11, 12).

Despite the wide adoption of bioluminescence-based assays, the limited thermal stability of the enzymes and substrates used to generate the assay signals continues to present a challenge. The reagents require long-term storage at ultra-cold temperatures and the performance of the reagents can degrade over time, adversely impacting sensitivity and assay reproducibility and limiting their use in point-of-care applications. Although significant efforts have been made to enhance the stability of bioluminescent reagents through genetic engineering and chemical modification (13, 14, 15, 16, 17, 18, 19), the components remain highly temperature dependent. Lyophilization can, in some cases, enhance reagent stability and reduce the need for cold storage, but lyophilization has significant drawbacks (20, 21, 22). The process is often variable and may result in reduced product yields. Solubility issues may prevent the coformulation of different reaction components in a single vial. Due to costs and process limitations, lyophilized samples are typically stored in relatively large, multi-use aliquots containing 1-10 mg/vial, creating the risk for cross-contamination and material waste. Alternative methods that can enable storage of bioluminescence reagents in stable, single-use quantities could improve data quality, reduce reagent waste, simplify laboratory workflows, and address many of the sustainability challenges associated with cold storage.

We previously described the application of Capillary Assisted Vitrification (CAV, previously referred to as Capillary Mediated Vitrification or CMV) as a platform technology for stabilizing enzymes, antibodies, and mRNA (23, 24, 25, 26). The CAV process uses a combination of inert small molecule stabilizers, a porous support material referred to as a scaffold, and rapid vacuum drying to transition biomolecules from a liquid state to a stable glassy state. The process is straightforward, can be completed in the lab using a benchtop instrument in under two hours, and requires little or no molecule specific optimization. Because the CAV technology can significantly enhance the thermostability of the stabilized molecules and can be applied to a broad range of materials with little or no formulation development, it provides a unique opportunity to create room-temperature stable reagents, reagent mixtures, and assay kits that are simple to use and eliminate the need for cold-chain storage and distribution. The use of these highly stable reagents may also reduce assay variability that results from loss of reagent activity over time. The stabilization technology can be easily deployed in any laboratory setting, making it a particularly attractive option for producing custom assay kits and stabilizing high-value custom reagents.

We previously characterized the synergistic interaction of CAV stabilizers, the scaffold, and the vacuum drying process and demonstrated how each of the components influences the CAV-process yield and the stability of the final product (24). Here, we evaluate the application of a variety of stabilized bioluminescent reagents in different types of real-world analytical workflows. We expand on our previous work by demonstrating both short and long-term stability of different luciferase enzymes and stabilization of the small molecule substrates, luciferin and furimazine. We establish that the reagents can be stabilized independently or co-formulated into a variety of ready-to-use assay master mixes using a common stabilization protocol. Finally, we evaluate the suitability of the CAV stabilized reagents in a wide range of *in vitro*, *in vivo,* and biochemical methods including cell viability assays, tumor imaging, and mix-and-read bioluminescent immunoassays. Overall, the results of this investigation indicate that CAV-stabilization is an attractive platform stabilization technology that enables long-term ambient storage while maintaining assay performance and offering significant advantages compared to traditional reagent storage paradigms.

## RESULTS

### Performance and Stability of Luciferase and Luciferin

Implementation of bioluminescence applications under real-world conditions require the ability to store luciferase and/or the substrates for periods of up to 6 months or more. If the reagents are shipped and stored under ambient conditions, they must also be capable of withstanding short-term exposure to significantly higher temperatures. Thus, we initiated both long-term stability studies of luciferase and luciferin at room temperature and a short-term study at 55 °C. At each time point, the activity of the CAV-stabilized reagents was evaluated by comparing the performance of the CAV-stabilized luciferase or luciferin to a reference preparation made from reagents that were stored frozen and thawed and prepared on the day of use (**Fig. 1**).

**Fig. 1.**
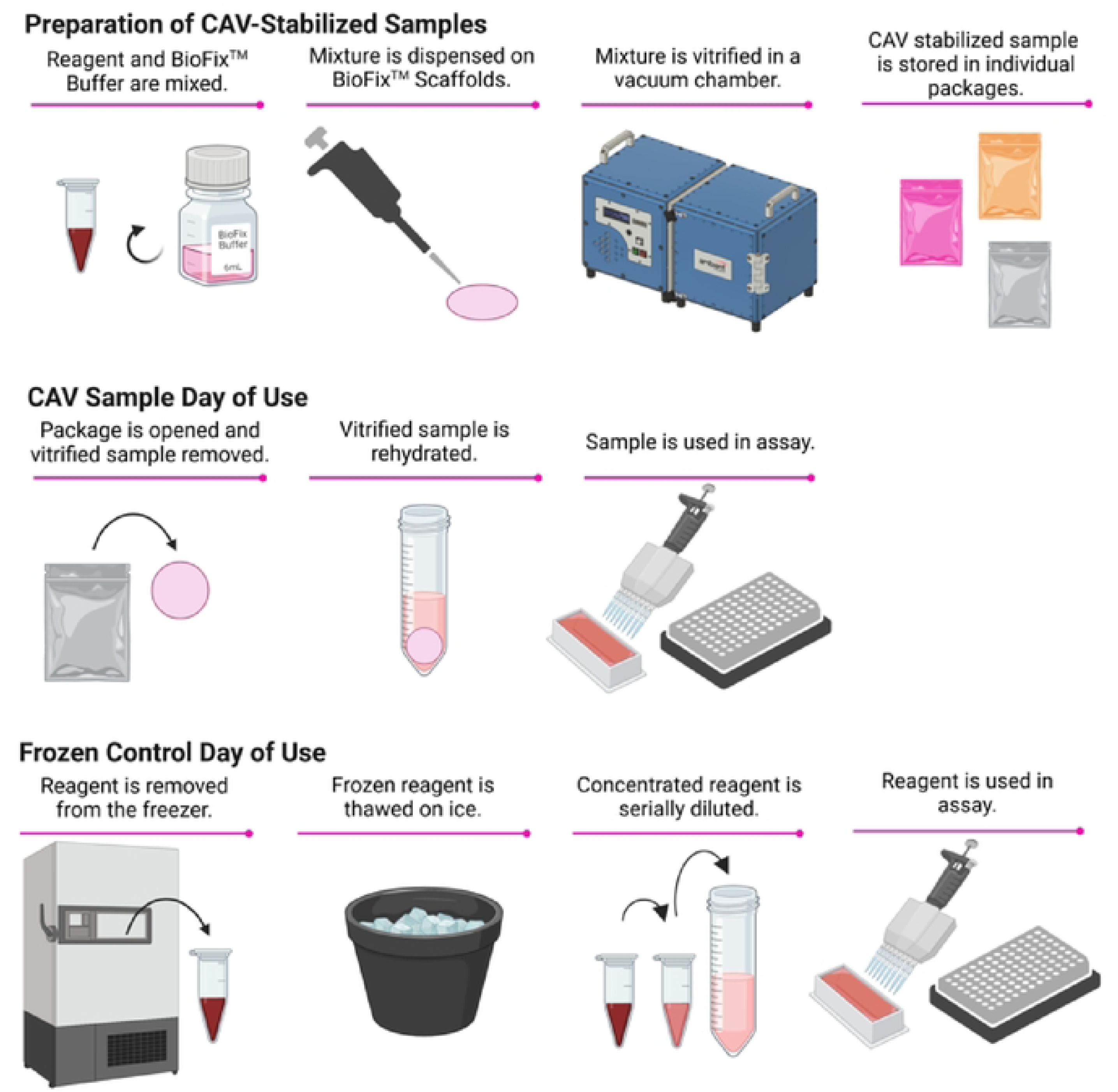
Schematic diagram of the CAV-stabilization process (top), rehydration and day-of-use protocol for CAV-stabilized samples (middle), and preparation of liquid controls from frozen reagent stocks (bottom).

At the start of the stability study, we evaluated the recovery of the two reagents immediately post-stabilization. As shown in **Fig. 2A**, the signals obtained for the CAV-stabilized luciferin were comparable to the frozen reference (p-value = 0.942). When the results for the CAV-stabilized luciferase were compared to the frozen luciferase reference material (**Fig. 2B**), a statistically significant difference between the CAV-stabilized luciferase was observed compared to the reference luciferase (p-value = 0.008). The CAV-stabilized material provided slightly higher signals and lower variability across replicate results compared to the frozen control. Despite the small difference in signal, the data indicates that the stabilization buffer does not interfere with the enzyme activity and that the process results in little or no loss of material.

**Fig. 2.**
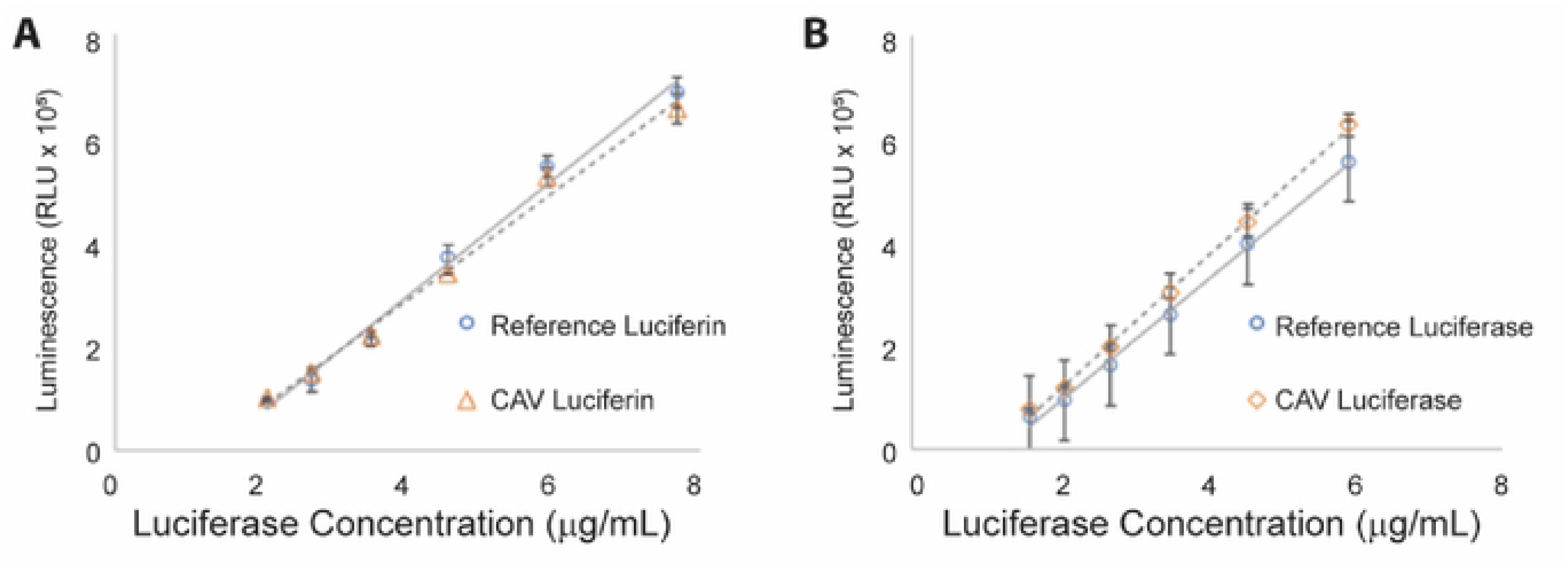
(**A**) Comparison of luminescent signal generated using CAV-stabilized luciferin (orange triangles) compared to a reference material from freshly thawed reagents (blue circles). (**B**) Comparison of luminescent signal generated using CAV-stabilized luciferase (orange diamonds) compared to a reference material from freshly thawed reagents (blue circles). Error bars show standard deviation with **p < 0.01 as determined by a paired two-sided t-test.

Next, aliquots of the CAV-stabilized form or matrix-matched, liquid controls were placed on stability. At the end of the study, stability data were evaluated graphically to estimate the degradation rates at each temperature. As shown in **Supplementary Figs. S1 and S2**, the CAV-stabilized materials show a linear trend when the natural log of activity is plotted versus time, indicating the degradation process can be described by a first-order reaction rate. According to the differential rate law, the slope of this line is equal to-k, where k is the reaction rate and the half-life (t_1/2_) can then be calculated using the equation:

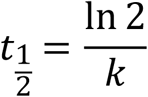

The t_1/2_ values are shown in **Table 1**. The t_1/2_ for the CAV-stabilized luciferin was estimated to be 39 months at room temperature and 2.1 months at 55 °C. The t_1/2_ values for CAV-stabilized luciferase was estimated to be 19 months at 55 °C. Although a linear trend was observed for luciferase stored in the CAV-form at 25 °C, the half-life could not be accurately measured. The activity of the CAV-stabilized luciferase was essentially flat across the nearly 2-year study, showing little or no downward trend and resulting in a correlation coefficient of 0.13. Even after 22 months of storage, the enzyme maintained 97% activity (**Supplementary Fig. S2**). The results suggest that the t_1/2_ will significantly exceed 2-years at room temperature.

**Table 1.** Comparison of the reaction rates and half-lives (t_1/2_) for luciferin and luciferase stored in liquid or post-CAV-stabilization at various temperatures.

| Reagent | Storage Temp. (°C) | Liquid Formulation | CAV Stabilized |
| --- | --- | --- | --- |
| | | $t_{1/2}$ (months) [95% CI] | |
| Luciferin | 25 | 0.1 [0.07, 0.4] | 39 [34, 46] |
|  | 55 | <1 day<sup>a</sup> | 2.1 [1.8, 2.4] |
| Luciferase | 25 | Highly variable activity at all timepoints | >2 years<sup>b</sup> |
|  | 55 | <1 day* | 19 [14,28] |

<sup>a</sup>No activity was detected after 1 day at 55 °C; $t_{1/2}$ could not be calculated and is therefore reported as <1 day.
<sup>b</sup>Due to small degree of change in luciferase activity during the study, the correlation between the natural log of activity and time is poor ( $R^2 = 0.13$ ) and the reaction rate and $t_{1/2}$ cannot be accurately determined for this condition.

With the exception of luciferin stored at room temperature (t_1/2_ = 0.1 months), we were unable to determine t_1/2_ for the liquid formulations. When stored as a liquid at 55 °C, both luciferase and luciferin lost all activity in less than 1 day. Test results for the luciferase formulation stored as a liquid at 25 °C were not reproducible between time points or between repeated experiments. Due to this variability, the performance of the liquid luciferase stored under this condition is considered unacceptable for an analytical reagent.

Overall, our stability evaluation confirms the following: 1) the CAV process significantly enhances the stability of both luciferase and luciferin, 2) the CAV-stabilized reagents can be stored long-term at room temperature, and 3), the CAV-stabilized reagents can be withstand short-term temperature excursions of up to 55 °C. Together, the data suggests that the CAV-stabilized reagents are suitable for ambient distribution and long-term storage.

To further evaluate the performance of the CAV-luciferase with respect to reagent consistency over time and between stabilization cycles, a scale-up study was performed. In each run, a total of 512 scaffolds were prepared and stabilized in a single stabilization cycle. A minimum of 6 scaffolds were tested immediately post-stabilization and again after the scaffolds had been stored for 9-months at room temperature. The mean activity across all four batches was 103% both post-stabilization and 103% after 9 months of storage (**Table 2**). The relative standard deviation (RSD) across scaffolds produced within a batch ranged from 1.9% to 7.8%. The RSD across the batches was 7.2% at time 0 and 5.2% after 9 months of storage. This data both demonstrates that the CAV-process is highly consistent and further confirms the observation that the stabilized reagents are stable for months or years at room temperature.

**Table 2.** Evaluation of CAV process consistency across four separate stabilization cycles. Comparison of luciferase activity on scaffolds immediately post-stabilization cycles and after 9 months of storage at room temperature.

| Full-Scale Run | Post Stabilization | 9M RT |
| --- | --- | --- |
|  | % Luciferase Activity (RSD) |  |
| Batch 1 | 99 (1.9%) | 97 (4.9%) |
| Batch 2 | 95 (2.8%) | 101 (3.4%) |
| Batch 3 | 112 (6.2%) | 107 (4.0%) |
| Batch 4 | 105 (7.8%) | 108 (5.8%) |
| Average | 103 (7.2%) | 103 (5.1%) |

RLU: Relative luminescence units

### Experimental Applications

#### Individually Stabilized Reagents

##### In Vivo Imaging

We next evaluated the suitability of the CAV-stabilized luciferin in an *in vivo* imaging application and compared results to those generated with the reference luciferin that was stored lyophilized or frozen. Luciferin was stabilized alone, rather than co-formulated with other reaction components. We quantified tumor burden using a 4T1 mouse mammary carcinoma model. Tumor size was monitored for up to 21 days after 4T1 tumor inoculation (**Fig. 3**), and the results obtained with bioluminescence imaging were compared to measurements made with calipers. We observed no statistical difference in the relationship between caliper measurements and bioluminescence signal of tumor growth between the luciferin groups (**Fig. 3A-C**). Although the CAV-stabilized luciferin experienced temperatures exceeding 65 °C during shipment, we found that both *in vivo* bioluminescence on day 16 post-inoculation and *ex vivo* tumor measurements were comparable in mean signal and that the CAV-stabilized luciferin exhibited a smaller variation in signal compared to the frozen control luciferin (**Fig. 3C-D**). This side-by-side analysis suggests that the stabilization process results in a functional and stable product equivalent to the current gold standard in bioluminescence imaging of tumors in mice.

**Fig. 3.**
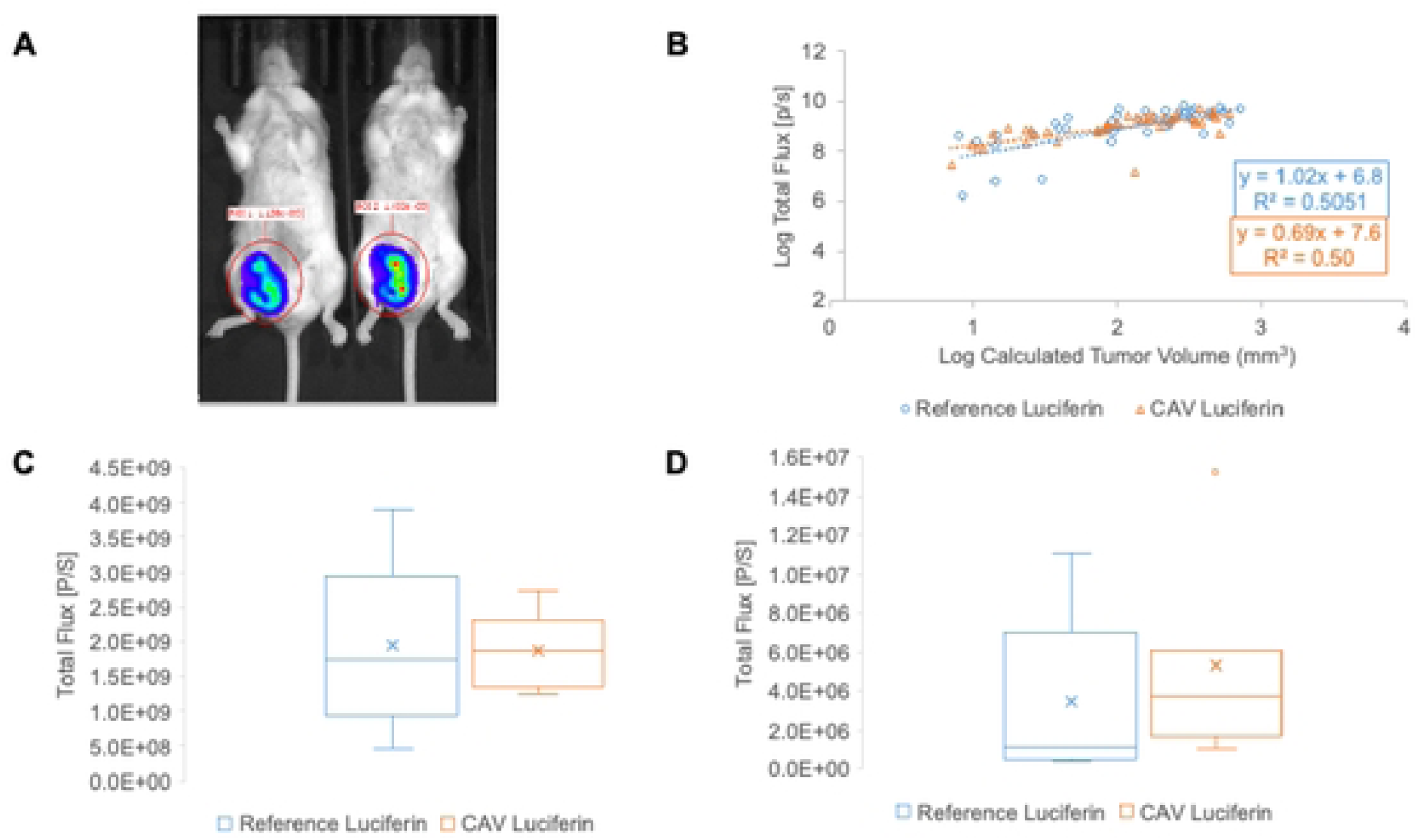
CAV-stabilized luciferin for quantitation of tumor burden and *in vivo* imaging. *In vivo* tumor imaging reveals consistent bioluminescence measurements between frozen reference (blue) or CAV-stabilized (orange) luciferin. (**A**) Mice implanted with 4T1 breast cancer cells in the #4 right mammary fat pad and imaged with the frozen reference luciferin (2 of 15 mice shown) (left) or CAV-stabilized luciferin (right). (**B**) Relationship between bioluminescence intensity and tumor volume based on caliper measurements. (**C**) Comparison of *in vivo* bioluminescence on day 16 post-inoculation (n = 7 reference, n = 8 CAV-stabilized). (**D**) Comparison of *ex vivo* tumor bioluminescence (n = 7 per group). The boxplots in C and D show the mean (x), median, the interquartile range (IQR), and ± 1.5 x IQR.

##### Reporter Gene Assays

Bioluminescence is frequently used to monitor the expression of reporter genes in cell culture applications. Cells containing the reporter gene are cultured, subjected to the test condition, and treated with luciferin or another substrate to produce a luminescent signal that is proportional to the level of reporter gene expression. To assess whether vitrified luciferin could be used in this type of application, luciferase-expressing 4T1 mouse breast cancer cells were prepared as a suspension culture and then serially diluted. **Fig. 4** shows that the frozen and CAV-vitrified luciferin signal are not statistically different.

**Fig. 4.**
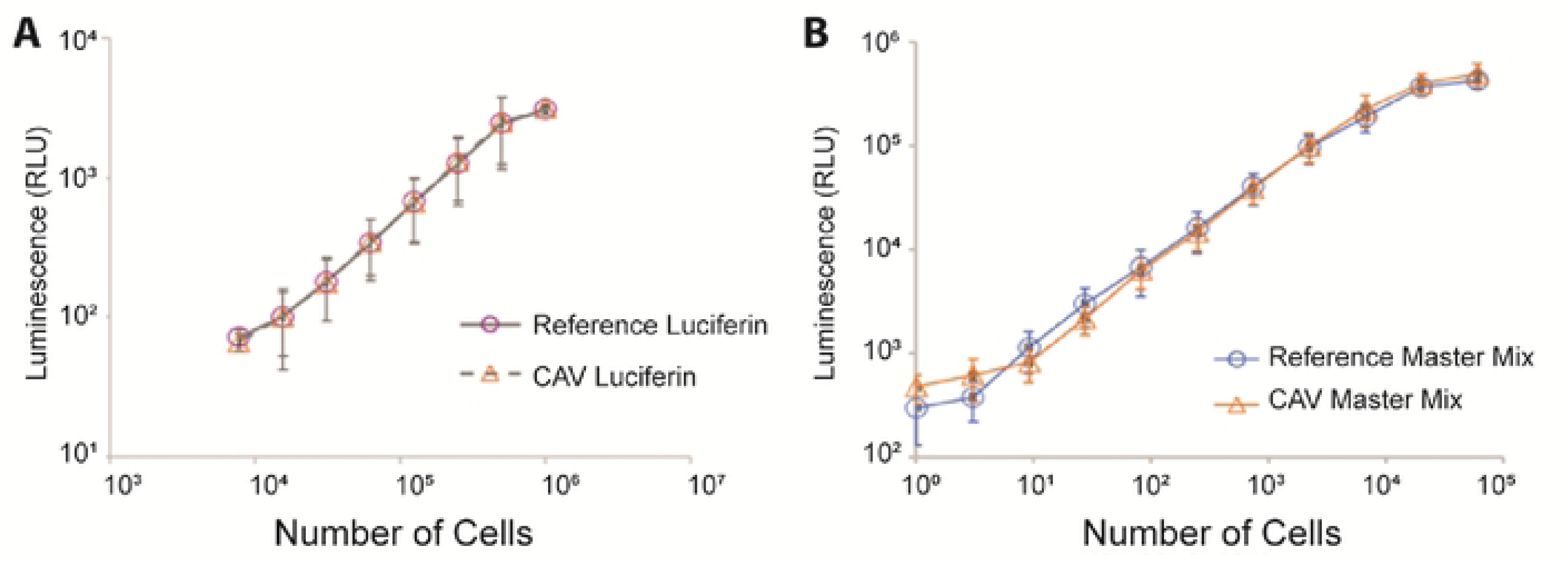
(**A**) Comparison of the signals generated using reference (blue circles) or CAV-stabilized luciferin (orange triangles) for quantification of gene expression in a 4T1 cell suspension culture (n = 3). (**B**) Comparison of cell viability data obtained using reference reagent master mix (blue circles) or CAV-stabilized master mix (orange triangles) (n = 4). Error bars represent standard deviation. Statistical significance determined by ANCOVA analysis (ns, nonsignificant).

##### Co-Formulated Master Mixes

As described above, many bioluminescent assays require multiple reagents that, under traditional storage paradigms, must be stored individually. On the day of use, each reagent must be located in the cold storage unit, thawed, diluted and co-formulated to produce a master mix. With the goal of developing thermally stable kits for bioluminescence testing, we prepared stabilized, assay mastermixes by co-formulating all of the reagents required to run a specific test method prior to stabilization. On the day of the test, a single scaffold was removed from the packaging, eluted, and immediately used in the appropriate assay. No further reagent preparation or dilutions steps were required prior to use. For each experiment, reagents stored in the traditional manner (lyophilized or frozen, with each reaction component stored in a separate vial), were used to prepare a control reagent mixture. The control or reference mixture was prepared fresh on the day of use.

##### Cell Viability Assessment

To test the performance of the CAV-stabilized master mix for cell viability applications, freshly cultured 4T1 cells were placed in a 96-well plate and diluted to 1 to 6E4 cells/well and treated with the reagent master mix. The results obtained with the frozen and CAV-stabilized mixture are visually indistinguishable and not statistically different (**Figure 4B**).

##### Microbial Contamination

To test for surface contamination (**Fig. 5**), slides were prepared by applying a buffer control (negative condition) or freshly cultured E. coli (positive condition). Swabs from the slides were placed into a vial containing either the traditional or CAV-stabilized reagent master mix. The slide was considered positive if the measured signal was above the detection limit. All samples were correctly identified as positive or negative using both the traditional and CAV-stabilized master mix (**Supplementary Table S1**).

**Fig. 5.**
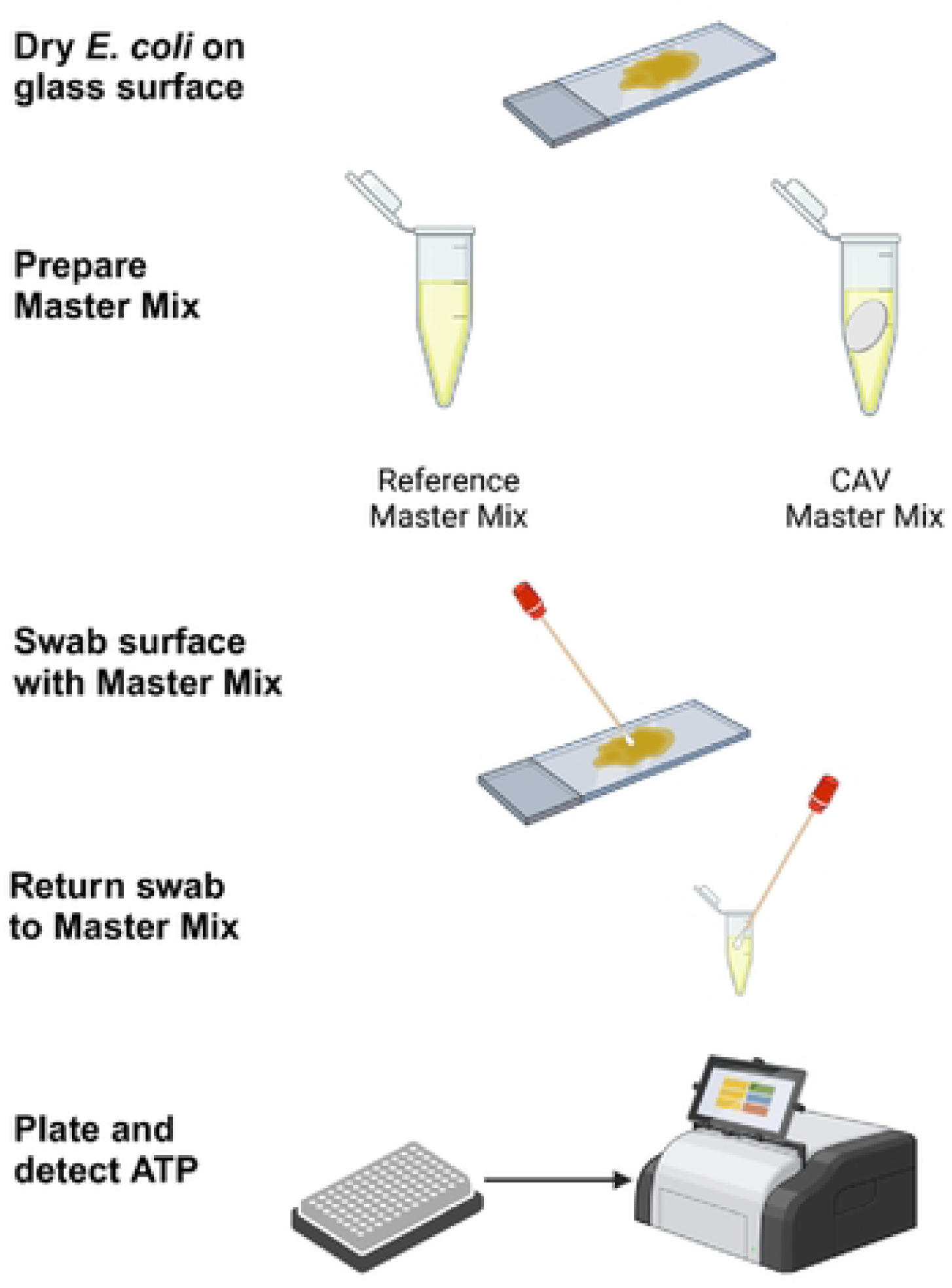
Schematic diagram of the microbial contamination assay. Glass slides were treated with E. coli or sterile buffer and dried. The slides were swabbed, and the swab was placed into a tube containing the reaction mixture prepared fresh or from CAV-stabilized reagents. The resulting luminescence was analyzed using a plate reader.

##### Homogeneous Immunoassay Kits

The Lumit® Flex technology is a homogeneous immunoassay that utilizes a ternary split NanoLuc® luciferase complementation system (27, 28, 29). Two small peptide tags (Peptide α and Peptide β) are chemically or genetically attached to analyte-specific antibodies or binding proteins. A detection polypeptide is then added, which remains inactive until it is brought into proximity to the peptide tags. When the ternary components assemble upon analyte binding, the fully reconstituted NanoLuc® enzyme catalyzes the furimazine substrate, generating a bright, stable bioluminescent signal. While each of the individual reaction components can typically be stored frozen or lyophilized, co-formulation of the components into a ready-to-use mixture can be challenging due to solubility and stability issues, particularly with the furimazine substrate.

Because the CAV-technology has been successfully used to stabilize both small and large molecule reagents, we explored whether CAV stabilization could be used to prepare an interleukin-6 (IL-6) Lumit® Flex master mix (**Fig. 6A**). The performance of the standard solution-based Lumit Flex IL-6 immunoassay was compared with a CAV-prepped Lumit Flex immunoassay in which all the assay components were co-stabilized. On the day of use, the master mix was reconstituted in PBS and an IL-6 standard curve generated and compared to standard curve generated with the standard, solution based reagents (**Fig. 6B**). Both assay formats show a dose-dependent increase in luminescent signal in response to increasing concentrations of recombinant human IL-6. The solution-based assay exhibits higher absolute signal intensity and a wider dynamic range, while the CAV-prepped immunoassay maintains consistent sensitivity and signal linearity despite a slightly lower overall luminescent output. Together, this reinforces the utility of the CAV process. If needed, the signal intensity could be improved further assay optimization or retitration of the reaction components.

**Fig. 6.**
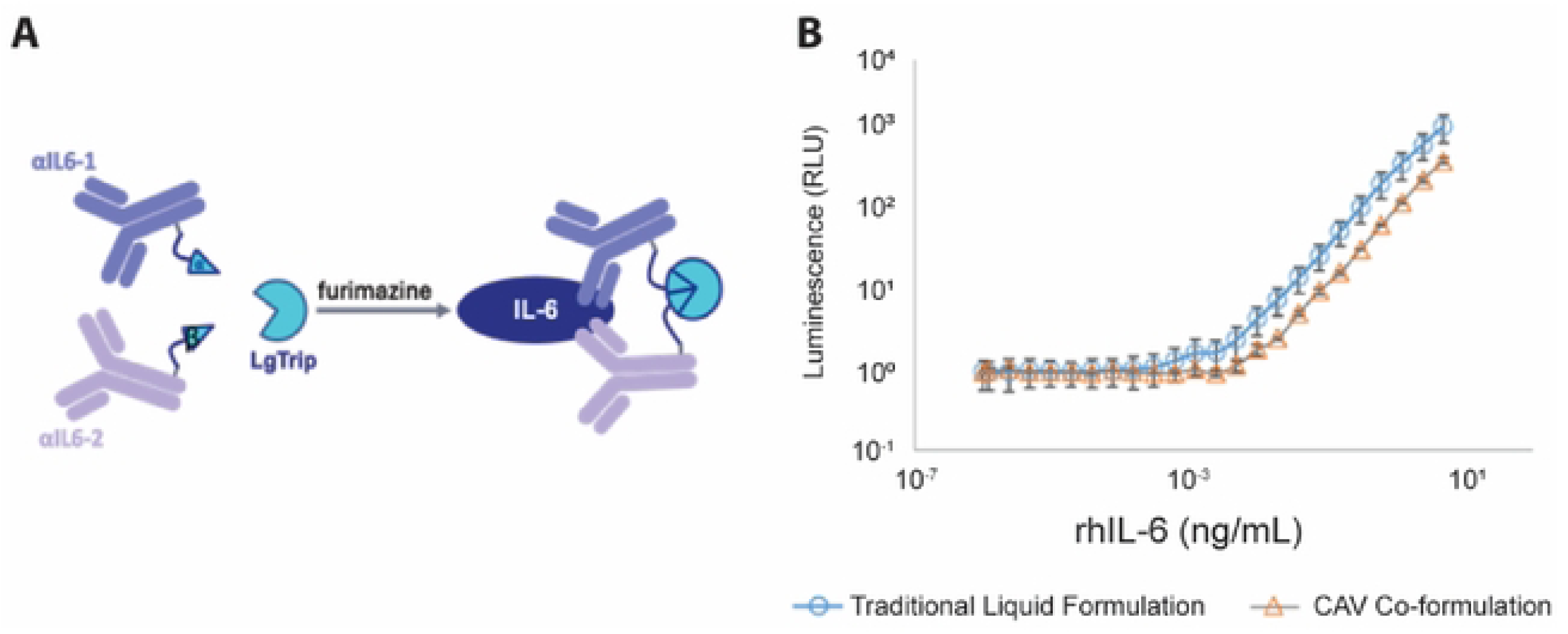
(**A**) Schematic diagram of the Lumit® Flex IL-6 Immunoassay. (**B**) Comparison of the performance of Lumit Flex immunoassay using traditional reagent formulation (blue circles) and a co-formulated master mix containing all assay components and stabilized using the CAV process (orange triangles). The CAV stabilized material shows a slight loss in absolute signal but comparable linearity and sensitivity. Error bars indicate the maximum and minimum signals observed across duplicate runs.

## DISCUSSION

We have established the effectiveness of CAV stabilization as a tool for enhancing the stability of a wide variety of bioluminescent reagents and the advantages of replacing traditional storage paradigms with newer technologies. In this study, a variety of luciferases, substrates, and master mixes were stabilized using the Ambient benchtop stabilization process. The reagents were all stabilized using the platform stabilization protocol. No optimization of the excipients or vacuum drying cycle was required, making the process easy to implement in any laboratory setting.

We showed that CAV stabilization significantly increases the t_1/2_ and useable shelf-life of both enzymes and small molecule substrates compared to liquid formulations. Two of the main challenges with current bioluminescence workflows include the need for ultra-cold reagent storage and assay reproducibility. Fluctuations in the assay performance from lot-to-lot or aliquot-to-aliquot variation in reagent performance can lead to data artifacts, reduced method sensitivity, and inaccurate conclusions. The results presented demonstrate the potential for CAV technology to address these crucial issues.

Across the range of assays evaluated, including both *in vitro* and *in vivo* analytical methods, the mean signals generated by the CAV-stabilized material were comparable to the traditional reagent format. One exception was the reduced signal intensity observed in the NanoLuc® assay, but this limitation could likely be overcome with additional assay optimization. Although we did not directly evaluate the impact on method precision, we anticipate that the improvement in the reagent stability over time will have long-term benefits on assay reportability and intermediate precision. The data shown in **Figs. 3 and 4** suggest that the CAV-stabilized reagents may also offer an opportunity to improve intra-assay variability. Freezing and lyophilization are both known to introduce heterogeneity and concentration gradients within small volume aliquots (30). This can adversely impact method performance resulting in variability between runs and/or between days. The CAV process therefore reduces the challenge of batch-to-batch variability.

Reformulation of analytical reagents is difficult as the formulation must be optimized not only to enhance the stability of the reagent and ensure consistent activity across individual aliquots and lots, but the stabilizers must not interfere with the assay signal. Our results indicate that the excipients used in the CAV-stabilization process are inert, do not interfere with spectroscopic analysis, and are safe to use in both cells and animals. Additionally, the same stabilization protocol can be applied with little or no reagent-specific customization. Thus, researchers can utilize this system to easily develop custom assays that will minimize reagent degradation and optimize assay performance over time.

Eliminating the need to store and ship reagents using cold or ultra-cold storage can have an immediate impact on laboratory carbon footprints and improve global access to research, clinical products, and testing supplies (31). The thermal stable microbial contamination kit would be a valuable tool for preventing infection in field hospitals and contamination in areas where maintaining proper food-hygiene may be challenging. Likewise, the CAV-stabilized Lumit Flex immunoassay demonstrates robust analyte detection while maintaining a simplified, shelf-stable format, making it highly suitable for point-of-care applications in low-resource environments. The CAV-stabilized immunoassay retains excellent sensitivity and dynamic range, ensuring reliable quantification of the IL-6 analyte, which is important in understanding multiple pathologies, response to therapy, and inflammation (32, 33, 34).

While the research described here focused solely on bioluminescence applications and specifically those that use the luciferase detection system, this platform technology is applicable to a wide range of applications. The stabilization process is fast and simple and can be executed by researchers with no specialized training. Thus, this platform can be used by researchers in the laboratory or in the field to prepare their own assay kits or stabilize their critical samples, thereby facilitating rapid, on-site diagnostic testing in decentralized settings such as rural clinics, field hospitals, and outbreak monitoring sites. This innovation enhances accessibility to sensitive, quantitative immunoassays where traditional laboratory infrastructure is limited.

## MATERIALS AND METHODS

### Sample Preparation

QuantiLum® Recombinant Luciferase, Beetle Luciferin, VivoGlow® Luciferin, and Lumit® Flex human IL-6 assay reagents were obtained from Promega Corp. (Madison, WI). Upon receipt, the luciferase was stored at-80 °C as per the manufacturer’s recommendations. The lyophilized beetle luciferin was reconstituted at 10 mg/mL in phosphate buffered saline (PBS; Cytiva, Logan UT)), aliquoted and stored at-80 °C. VivoGlo Luciferin was reconstituted in a 1:1 mixture of PBS and BioFix^TM^ stabilization buffer, and aliquots stored at-20 °C.

CAV stabilized samples were prepared in the lab using the Ambient Stabilizer (Ambient Biosciences, Ann Arbor MI). The CAV process was described previously (23), and a schematic of the process is provided in **Fig. 1**. Briefly, the reagent of interest is prepared at a 2X concentration and mixed with and equal volume of BioFix™ or BioFix+™ stabilization buffer (Ambient Biosciences, Ann Arbor MI). The formulated material is dispensed onto a porous material referred to as the scaffold. The quantity of reagent is determined based on the assay requirements and generally each scaffold is designed to hold enough reagent for a single assay run.

The BioFix™ buffer is a commercially-available reagent that contains a mixture of saccharides and other inert small molecule stabilizers. The buffer serves two key roles in the stabilization process, facilitating the formulation of an amorphous glass during drying and replacing the hydrogen bonds that are lost when water is removed from the aqueous system. The BioFix+™ stabilizer containers the same stabilizers but also includes a carrier protein designed to prevent non-specific binding of proteins to surfaces. The BioFix+™ is recommended when working with low concentration protein reagents but can be used for other reagents including small molecules.

For this study, scaffolds were selected from a catalog of commercially available option that differ primarily in terms of the quantity of material that can be applied and stabilized on each scaffold. For each experiment, the smallest scaffold that would hold a suitable volume of the stock reagent was selected. Each scaffold has a small hold up volume, or volume of liquid that cannot easily be extracted. Using the smallest possible scaffold reduces loss and results in optimal yields. The VivoGlo luciferin, luciferase, and microbial contamination kits were therefore stabilized using BioFix™ 175 Scaffolds (Ambient Biosciences, Ann Arbor, MI). All other materials were stabilized using BioFix™ Classic Scaffolds (Ambient Biosciences, Ann Arbor, MI).

On the day of use, the CAV-stabilized samples were rehydrated by incubating the vitrified scaffolds in elution buffer. The choice of elution buffer was determined separately for each assay based on the assay requirements. Incubation times of 1 minute or less were sufficient to fully rehydrate the CAV-stabilized material. Unless otherwise noted, the eluted material was used in the assay with no further manipulation. **Fig. 1** provides a schematic of the rehydration process and for comparison pursues, the workflow used to prepare liquid controls.

### Luciferase and Luciferin Activity Assay

Luciferase and luciferin activity were measured using a microtiter plate-based assay. A luciferin reaction mixture containing 1.5 mM beetle luciferin, 4.5 mM magnesium sulfate (Sigma, St. Louis, MO), and 0.5 mM ATP (Sigma, St. Louis MO) was prepared in 25 mM EPPS buffer (Sigma. St. Louis MO), pH 7.8. Separately, serial dilutions of luciferase ranging in concentration from 0 to 23 µg/mL were prepared. To initiate the assay, a 50 µL aliquot of each luciferase dilution was placed in a 96-well flat-bottom plate with a clear bottom and black walls (Corning, Corning NY) and mixed with a 50 µL aliquot of the luciferin reaction mixture. The plates were incubated for 8 minutes after the addition of all reagents and luminescence was measured using a Synergy H1 microplate reader (BioTek, Winooski, VT) and the endpoint luminescence protocol.

The activity of the test articles was determined by comparing the response of the test article with a reference standard. The reference standard was made using a luciferin mixture and luciferase dilutions that were freshly prepared from frozen stocks on the day of use. The relative activity of the test article was calculated by direct interpolation for each sample dilution that fell within the linear range of the assay and averaging across the dilution adjusted results.

The test articles consisted of one of the following: 1) CAV-stabilized luciferase, 2) luciferase stored in a liquid formulation, 3) CAV-stabilized luciferin, or 4) luciferin stored in a liquid formulation. When luciferase was tested in either the CAV-stabilized or liquid formulation, freshly thawed and diluted luciferin was used. Likewise, when testing luciferin test articles, a freshly prepared luciferase solution was used.

### Luciferase Stability Assessments

To evaluate the stability of CAV-stabilized luciferase and luciferin, the reagents were initially stabilized individually. CAV-stabilized luciferin samples were prepared so that each scaffold contained 239 µg luciferin in BioFix+™ buffer. At the time of testing, the luciferin scaffolds were eluted with 0.5 mL of a solution containing 4.5 mM magnesium sulfate, 0.5 mM ATP in 25 mM EPPS buffer, pH 7.8. A liquid control was also prepared. The composition of the liquid controls was identical to that of the CAV-stabilized samples post elution with the exception that the liquid controls did not include the BioFix^TM^ stabilization buffer.

CAV-stabilized luciferase samples were prepared so that each scaffold contained 92 µg of luciferase in BioFix+™ buffer. At the time of the test, the luciferase scaffolds were eluted with 4 mL of PBS, producing a solution containing 23 µg/ml luciferase which was then used to prepare serial dilutions for testing. Again, a liquid control was prepared that matched the composition of the luciferase eluted from the BioFix™ scaffolds, but did not include the BioFix+™ stabilization buffer.

The CAV-stabilized samples and liquid controls were stored on the bench (room temperature condition) or in a temperature-controlled incubator set at 55 °C. For the CAV-stabilized samples, three scaffolds were removed from the incubator at each time point. Triplicate scaffolds were eluted and independently tested, generating three test results per time point. For the liquid controls, a single aliquot was removed at each time point. The liquid control was sampled and tested three times.

### Tumor Imaging in Mice

All animal experiments were carried out according to a protocol approved by the Vanderbilt University Institutional Animal Care and Use Committee. Bioluminescence was evaluated in mice using luciferase-expressing 4T1 cell lines. The CAV-stabilized luciferin samples used in this experiment contained 9 mg of VivoGlo® Luciferin per scaffold. A reference solution was prepared using lyophilized VivoGlo® luciferin reconstituted in PBS. Materials for the tumor imaging studies were shipped to the testing site ambiently via ground transportation and stored at room temperature until use. Ground transportation required 6 days, and during this time the material was exposed to temperatures ranging from 71-107 °F (21-42 °C). The corresponding liquid controls were shipped on dry ice.

Seven-week-old BALB/c female mice (Charles River Laboratories) were implanted with 5×10^4^ 4T1-luciferase-expressing cells, suspended in 50 μl of PBS, in the #4 right mammary gland. On the day of use, scaffolds were eluted in 1 mL per scaffold resulting in a final post-elution concentration of 9 mg/mL. Excess reference solution was stored as frozen aliquots between imaging sessions. Bioluminescence imaging 2x/week followed 30 min after a 150 mg/kg i.p. injection of CAV-stabilized luciferin or reference luciferin (n = 7 reference, n = 8 CAV-stabilized). To minimize pain and distress, discomfort produced from tumor implantation was controlled using inhaled isoflurane anesthetic by nose cone (2% isoflurane in 100% O_2_, flow rate 1 L/min). During imaging, each subject was anesthetized on a heated platform using isoflurane delivered via an induction chamber and 2% isoflurane lines. Imaging is non-invasive, and anesthesia was introduced only to keep the subject still during the experiments. We monitored all mice for potential signs of distress over the course of the experiment. Tumors were imaged using an IVIS Lumina Series III microscope (Caliper LifeSciences). Tumor length and width were measured using digital calipers 2x/week up to 21 days post-inoculation or when humane endpoint (tumor volume>1,000 mm^3^) was reached. Tumor volume was calculated using the formula: Volume = (D_1_^2^x D_2_)/2, where D_1_ is the minimum diameter and D_2_ is the maximum diameter. Tumors reached an average of 650 ± 238 mm^3^, and one mouse died before meeting the criteria for euthanasia.

### 4T1 Reporter Gene Expression

A luciferase-expressing mouse mammary carcinoma cell line 4T1/Puro Dual Labeled Stable Cell Line was obtained from GeneCopoeia (SL020). The cell line is stably transfected with both a firefly luciferase gene and eGFP gene. Cells were propagated and maintained in RPMI selection medium (BioWhittaker/Lonza, Walkersville MD and Gibco, Grand Island NY) containing 1% GlutaMAX (Gibco, Grand Island NY), 1% Ab/Am, puromycin (Gibco, Grand Island NY) and 10% FBS (Gibco, Grand Island NY). Cells were cultured at 37 °C in a 5% CO_2_ atmosphere.

On the day of testing, cells were suspended in medium containing 450 µg/mL luciferin and titrated from 10^3^ to 10^6^ cells per well in an opaque white 96-well cell culture plate (Corning, Corning NY). Prior to use, the luciferin was either stored frozen and thawed and diluted in media on the day-of-use (reference solutions) or in the CAV stabilized form. Each CAV-stabilized aliquot contained 2.7 mg of beetle luciferin and was eluted in 6 mL of media. After a 15-minute exposure, bioluminescence intensity was read using a Synergy H1 microplate reader. The experiment was carried out on each of 3 days for n=3 replicate runs. The relative activity for the CAV-stabilized master mix was determined by calculating the average of all dilutions that fell within the linear range of the assay and reporting that result as a percentage relative to the reference standard. The data was background corrected by subtracting the signals obtained for a media only control. ANCOVA analysis was performed to test for statistical differences in the data set.

### CAV Master Mix for Cell Viability Assessment

Bioluminescence was evaluated in CRL-2539 4T1 cells (ATCC, Manassas VA) to evaluate whether luciferase and luciferin could be co-formulated into an easy-to-use assay master mix or kit for cell viability testing. This cell line does not contain a luciferase reporter gene. Cells were titrated from 1-6×10^4^ cells/well in an opaque, white 96-well cell culture plate, centrifuged, and washed with Dulbecco’s Phosphate Buffered Saline (DPBS, Lonza, Walkersville MD). 20 µL of lysis buffer containing 25 mM trisphosphate (pH 7.8), 2 mM dithiothreitol, 2 mM EDTA, 10% glycerol, 1% Triton X-100 was added to each well and the plate was incubated on a shaker for 20 minutes. Scaffolds containing 4.4 µg luciferase and 840 µg luciferin co-vitrified in BioFix+ stabilization buffer were prepared. Six scaffolds were eluted together in 5.28 mL buffer containing 4.5 mM magnesium sulfate and 25 mM EPPS, pH 7.8, for a final concentration of 5 µg/mL luciferase and 3 mM luciferin. A reference standard was prepared freshly thawed luciferase and luciferin diluted to 5 µg/mL luciferase and 3 mM luciferin in buffer containing 4.5 mM magnesium sulfate, 0.28% bovine serum albumin (BSA), 25 mM EPPS, pH 7.8. Subsequently, plate wells were filled with 100 µL master mix with sample or standard. After a 15-minute exposure, bioluminescence intensity was read using a Synergy H1 microplate reader. The relative activity for the CAV-stabilized master mix was determined by calculating the average across all dilutions that fell within the linear range of the assay and reporting that result as a percentage relative to the reference standard. ANCOVA analysis was performed to test for statistical differences in the data set.

### CAV Master Mix for Microbial Contamination Testing

A microbial contamination kit was prepared by vitrifying 0.875 µg luciferase, 139 µg luciferin, 9 µg DTT, and 0.035 µg Triton-X in BioFix+ stabilization buffer on BioFix 175 scaffolds. This master mix contained the enzyme, substrate and all co-factors required for testing in a single CAV-stabilized preparation. Sterile broth or broth containing 1*10^5^ e. coli cells was applied to sterile microscope slides and allowed to dry for 2 hours at room temperature. On the day of the test, a liquid reference control mixture was prepared from 4.16 µg/mL luciferase, 2.5 mM luciferin, 0.333 mM DTT, 2% glycerol, 0.2% triton, 3.5 mM MgSO_4_ in 25 mM Tris buffer. CAV scaffolds were eluted in 0.5 mL buffer containing 2% glycerol, 3.5 mM MgSO_4_ in 25 mM Tris. A FLOQSwab (Copan Cat #534CS01) was soaked in 0.5 mL reaction mixture, used to separately swab the surface of each glass slide, and then returned to a tube containing the reference master mix prepared from frozen stocks or from the CAV-stabilized master mix. A 100 µL quantity of the reaction mixture was then transferred to a 96-well plate and read for luminescence in a Synergy H1 plate reader. The microbial contamination assay format is shown schematically in **Fig. 5**.

### Co-formulated Lumit® Flex IL-6 Immunoassay

Anti-IL-6 antibody clones 5IL6 and 505E (Thermo Fisher) were chemically conjugated with the Lumit® Flex alpha and beta peptide (Promega Corp), respectively, and optimized for working concentration as previously reported(27). Briefly, each antibody was incubated with an amine-reactive form of the peptide labels for 1 hour at room temperature, quenched with 2M Tris buffer, and then diluted in 80% glycerol for storage. A solution-based assay was established using a master mix containing 2µM Lumit® Flex Detection Protein, 5IL-6-Lumit® Flex Peptide Alpha labeled antibody, 505E-Lumit® Flex Peptide Beta labeled antibody, and 10µM of NanoGlo Live Cell Substrate (Promega Corp). This master mix/coformulation was mixed with recombinant IL-6 (R&D Systems) titration prepared in assay buffer. After 1.5 h, the luminescence was read on a GloMax Discover Luminometer (Promega Corp).

The CAV-stabilization of the Lumit Flex IL-6 immunoassay was performed using the BioFix+ buffer. Post-stabilization, scaffolds were eluted with 500uL of PBS + 0.01% BSA to collect the assay components, which were then added to a non-binding white 96 well plate (Corning) containing a dilution of rhIL-6.

## STATEMENTS AND DECLARATIONS

### Ethics Approval and Animal Welfare

All animal experiments were carried out according to a protocol approved by the Vanderbilt University Institutional Animal Care and Use Committee and adhered to all relevant regulations and guidelines.

### Author Contributions

All authors contributed to this study. The study was conceived and supervised by M. Dart, M. Rafat, L. Bronsart, and M. Shank-Retzlaff. Material and equipment preparation, data collection, and analysis were performed by S. Radford, Y. Peris-Taverner, M. Dibble, K. Corn, T. Zhu, S. Martello, M. Mayeaux, A. Ladd, S. Renu, T. Chunduri, and A. Jadhav. The first draft of the manuscript was written by M. Shank-Retzlaff. All authors read and approved the final version of the manuscript.

### Data Availability

The data that support the findings of this study are available from the corresponding author upon reasonable request.

## Acknowledgements

The authors thank Dr. Pravansu Mohanty for technical support and Dr. Craig L. Duvall for IVIS use. Schematics were prepared using BioRender.

## REFERENCES

1. Deshpande SS. Principles and applications of luminescence spectroscopy. Crit Rev Food Sci Nutr. 2001;41(3):155–224.

2. Dunuweera AN, Dunuweera SP, Ranganathan K. A Comprehensive Exploration of Bioluminescence Systems, Mechanisms, and Advanced Assays for Versatile Applications. Biochem Res Int. 2024;2024:8273237.

3. Kulmala S, Suomi J. Current status of modern analytical luminescence methods. Anal Chim Acta. 2003;500(1-2):21–69.

4. Roda A, Pasini P, Mirasoli M, Michelini E, Guardigli M. Biotechnological applications of bioluminescence and chemiluminescence. Trends Biotechnol. 2004;22(6):295–303.

5. Syed AJ, Anderson JC. Applications of bioluminescence in biotechnology and beyond. Chem Soc Rev. 2021;50(9):5668–705.

6. Gleneadie HJ, Dimond A, Fisher AG. Harnessing bioluminescence for drug discovery and epigenetic research. Front Drug Discov. 2023;3.

7. Mali SB. Bioluminescence in cancer research - Applications and challenges. Oral Oncology Reports. 2024;9:100127.

8. Coleman SM, McGregor A. A bright future for bioluminescent imaging in viral research. Future Virol. 2015;10(2):169–83.

9. Baljinnyam B, Ronzetti M, Simeonov A. Advances in luminescence-based technologies for drug discovery. Expert Opin Drug Discov. 2023;18(1):25–35.

10. Nante N, Ceriale E, Messina G, Lenzi D, Manzi P. Effectiveness of ATP bioluminescence to assess hospital cleaning: a review. J Prev Med Hyg. 2017;58(2):E177–E83.

11. Whitehead KA, Smith LA, Verran J. The detection of food soils and cells on stainless steel using industrial methods: UV illumination and ATP bioluminescence. Int J Food Microbiol. 2008;127(1-2):121–8.

12. Deininger RA, Lee J. Rapid detection of bacteria in drinking water. Nato Sci S Ss Iv Ear. 2005:71–8.

13. Mehrabi M, Hosseinkhani S, Ghobadi S. Stabilization of firefly luciferase against thermal stress by osmolytes. Int J Biol Macromol. 2008;43(2):187–91.

14. Ganjalikhany MR, Ranjbar B, Hosseinkhani S, Khalifeh K, Hassani L. Roles of trehalose and magnesium sulfate on structural and functional stability of firefly luciferase. J Mol Catal B-Enzym. 2010;62(2):127–32.

15. Koksharov MI, Ugarova NN. Thermostabilization of firefly luciferase by in vivo directed evolution. Protein Eng Des Sel. 2011;24(11):835–44.

16. Lohrasbi-Nejad A, Torkzadeh-Mahani M, Hosseinkhani S. Hydrophobin-1 promotes thermostability of firefly luciferase. FEBS J. 2016;283(13):2494–507.

17. Jazayeri FS, Amininasab M, Hosseinkhani S. Structural and dynamical insight into thermally induced functional inactivation of firefly luciferase. PloS one. 2017;12(7):e0180667.

18. Pozzo T, Akter F, Nomura Y, Louie AY, Yokobayashi Y. Firefly Luciferase Mutant with Enhanced Activity and Thermostability. ACS Omega. 2018;3(3):2628–33.

19. Shi C, Killoran MP, Hall MP, Otto P, Wood MG, Strauss E, et al. 5,5-Dialkylluciferins are thermal stable substrates for bioluminescence-based detection systems. PloS one. 2020;15(12):e0243747.

20. Williams NA, Polli GP. The lyophilization of pharmaceuticals: a literature review. J Parenter Sci Technol. 1984;38(2):48–59.

21. Bhatnagar BT, S. Advances in Freeze Drying of Biologics and Future Challenges and Opportunities. Drying Technologies for Biotechnology and Pharmaceutical Applications 2020. p. 137–77.

22. Mirasol F. Lyophilization Presents Complex Challenges. Biopharm Int. 2020;33(1):22–4.

23. Shank-Retzlaff M, Taverner YP, Joshi P, Renu S, Chitikela A, Koneru A, et al. Capillary-Mediated Vitrification: A Novel Approach for Improving Thermal Stability of Enzymes and Proteins. J Pharm Sci. 2022;111(8):2280–7.

24. Renu S, Shank-Retzlaff M, Sharpe J, Bronsart L, Mohanty P. Capillary-Mediated Vitrification: Preservation of mRNA at Elevated Temperatures. AAPS J. 2022;24(4):75.

25. Amle S, Radford S, Wang Z, Bronsart L, Mohanty P, Renu S, et al. Use of capillary-mediated vitrification to produce thermostable, single-use antibody conjugates as immunoassay reagents. J Immunol Methods. 2023;516:113460.

26. Shank-Retzlaff M, Abdeen SJ, Bronsart L, Cieslak AN, Cruse JK, Kinne AS, et al. Capillary mediated vitrification is a novel technique that enables storage of antibody critical reagents at ambient temperature: Impact on binding, structure, and laboratory sustainability. J Pharm Biomed Anal. 2024;251:116409.

27. Kincaid VA, Wang H, Sondgeroth CA, Torio EA, Ressler VT, Fitzgerald C, et al. Simple, Rapid Chemical Labeling and Screening of Antibodies with Luminescent Peptides. ACS Chem Biol. 2022;17(8):2179–87.

28. Torio EA, Ressler VT, Kincaid VA, Hurst R, Hall MP, Encell LP, et al. Development of a rapid, simple, and sensitive point-of-care technology platform utilizing ternary NanoLuc. Front Microbiol. 2022;13:970233.

29. Hall MP, Unch J, Binkowski BF, Valley MP, Butler BL, Wood MG, et al. Engineered luciferase reporter from a deep sea shrimp utilizing a novel imidazopyrazinone substrate. ACS Chem Biol. 2012;7(11):1848–57.

30. Deck LT, Ferru N, Kosir A, Mazzotti M. Visualizing and Understanding Batch Heterogeneity During Freeze-Drying Using Shelf-Scale Infrared Thermography. Ind Eng Chem Res. 2024;63(38):16335–46.

31. Xu Q, Deng B, Wang YP, Liu WS, Chen G. Small, affordable, ultra-low-temperature vapor-compression and thermoelectric hybrid freezer for clinical applications. Cell Rep Phys Sci. 2023;4(12).

32. Brabek J, Jakubek M, Vellieux F, Novotny J, Kolar M, Lacina L, et al. Interleukin-6: Molecule in the Intersection of Cancer, Ageing and COVID-19. Int J Mol Sci. 2020;21(21).

33. Lin SW, Shen CF, Liu CC, Cheng CM. A Paper-Based IL-6 Test Strip Coupled With a Spectrum-Based Optical Reader for Differentiating Influenza Severity in Children. Front Bioeng Biotechnol. 2021;9:752681.

34. Grebenciucova E, VanHaerents S. Interleukin 6: at the interface of human health and disease. Front Immunol. 2023;14:1255533.

